# Asexual lineages of a cosmopolitan pest aphid are associated with different climatic niches

**DOI:** 10.1101/2023.08.10.552807

**Authors:** Martin Godefroid, Christine N Meynard, Anne-Laure Clamens, Megan Popkin, Emmanuelle Jousselin

## Abstract

Asexual lineages often exhibit broader distributions and can thrive in extreme habitats compared to their sexual counterparts. Two hypotheses have been proposed to explain this phenomenon. The general-purpose genotype model posits that selection favours a few versatile asexual genotypes with wide environmental tolerance, enabling their long-term persistence across diverse conditions. Conversely, the frozen niche variation model suggests that selection favours specialised genotypes with minimal niche overlap among them and their sexual relatives, potentially leading to competition-driven exclusion of both sexual and asexual relatives. To investigate these hypotheses, we examined ecological niche differentiation among six globally distributed obligate asexual lineages of the cosmopolitan aphid pest, *Brachycaudus helichrysi*. We initially investigated the presence of different endosymbionts across clones, as endosymbionts play a major role in aphid niche differentiation. Subsequently, we conducted multivariate analyses to explore climatic niche divergence among clones. We provide evidence for climatic niche specialisation in asexual lineages, which is not related to variations in endosymbiont associations. Therefore, despite their apparent global distribution, *B. helichrysi* clones exhibit characteristics of specialised genotypes, which aligns with the frozen niche variation hypothesis. This study represents the first comprehensive evidence of climatic adaptation in aphid superclones, providing novel insights into their global distribution.

## BACKGROUND

Understanding the ecological differentiation processes underlying the transition from sexuality to asexuality has been a longstanding conundrum for evolutionary biologists [1]. This inquiry has primarily focused on explaining why asexual lineages of many species exhibit wide geographic distributions and surpass their sexual counterparts in abundance, particularly in islands, disturbed habitats, and, more generally, environments with extreme conditions (a pattern known as “geographical parthenogenesis”) [2,3]. Two theoretical models have been proposed to explain the emergence, spread, and long-term persistence of asexual lineages, each generating distinct predictions about their ecological niches. The general-purpose genotype (GPG) hypothesis, originally proposed to explain the ubiquity of weedy plants [4], predicts the emergence of asexual lineages with broader environmental tolerances than their sexual relatives [5,6]. According to the GPG model, these asexual generalist lineages will exhibit great plasticity, enabling their persistence across extensive geographical areas and in rapidly changing habitats. Another theoretical model, referred to as the ‘frozen niche variation’ (FNV) hypothesis, which shares similarities with the “tangled bank model” [7], predicts that selection will favour specialised asexual genotypes with minimal niche overlap among closely related asexual and sexual lineages [8,9]. The rationale behind the FNV model is that repeated transitions to asexuality result in the selection of locally adapted genotypes from sexually reproducing populations. These genotypes efficiently use underutilised resources, especially in spatially heterogeneous environments [8–10]. In other words, the selection of asexual clones *freezes* ecologically-relevant genetic variation that already exists [9].

The respective predictions of both hypotheses have been explored in a wide range of organisms. Support for both, as well as inconclusive results, have been reported [10]. Two approaches have been employed in the study of asexual lineages’ ecological niches: (1) experimental approaches that compare the fitness of asexual clones between each other and/ or with their sexual relatives along environmental gradients under controlled conditions; (2) phylogeographic approaches that compare the distribution and realised ecological niches of different asexual clones and sometimes their sexual relatives [10]. While comparing the environmental response of different asexual lineages seems to be the most direct method for testing the predictions of FNV *vs* GPG, this approach remains relatively rare as it is time-consuming and requires being able to breed different lineages under laboratory conditions, which is not feasible for all organisms. Phylogeographic studies overcome these difficulties; however, they often come with their own set of limitations. For instance, in early studies that use distributional data, clonal lineages were sometimes considered as a single unit and then compared to their sexual relatives, without distinguishing whether clonal populations were composed of multiple independent clonal lineages or a single clonal lineage [11–13]. This could be due to the challenges associated with assigning individuals to different asexual lineages before genetic studies became standard practice in ecological research. This weakness prevents accurate investigation of the niche breadths of existing clones and may yield misleading patterns where asexual lineages appear to have broad ecological tolerances. In addition, many studies failed to utilise environmental data extracted at sampling sites [14], which prevents addressing whether geographically widespread asexual clones - that could be first seen as a generalist genotypes – actually occupy a narrow but widely available realised niche. Conversely, when clones tend to be geographically restricted to a few sites [15], it is challenging to ascertain whether this narrow distribution reflects true ecological specialisation, recent emergence from their sexual relatives, or simply a lack of opportunities for dispersal. To overcome all these caveats, phylogeographic approaches should ideally incorporate modern ecological niche approaches on a cosmopolitan species characterised by well-structured genetic populations, wherein asexual lineages are widely distributed.

The study of the leaf curl plum aphid, *Brachycaudus helichrysi* (Kaltenbach 1843) (Hemiptera, Aphididae) fits some of these prerequisites. This worldwide pest species has been recorded in more than 60 countries throughout the Palearctic [16] and is also present in the Southern Hemisphere [17]. As most aphid species, the typical life cycle of *B. helichrysi* is cyclical parthenogenesis. This involves a sexual reproduction phase occurring in autumn on *Prunus* species (mainly *Prunus domestica*), resulting in the production of overwintering eggs. Subsequently, an asexual reproduction phase occurs in spring and summer on a large range of herbaceous plants [17,18]. Recent genetic studies have unveiled that *B. helichrysi* is, in fact, a species complex composed of three distinct lineages referred to as H1, H2 and H3 [19–22]. Among these lineages, H2 primarily comprises populations that reproduce asexually all year round on herbaceous plants in areas with mild winters [19], enabling the persistence of parthenogenesis [23]. So far, few sexual populations have been identified, geographically restricted to Northern India and Central Asia and reproducing sexually on peach trees (*Prunus persica*) [19,20]. Within the H2 lineage, eight asexual lineages have been described using microsatellite markers and experimental studies have confirmed irreversible loss of sexuality in one of these clonal lineages [19]. Some of these clones have been found across multiple continents and over 15-year long stretches [19,20], thereby meeting the definition of aphid superclones [24,25]. Published analyses revealed that these clonal lineages do not exhibit significant differences in their associations with host plants [19]. Based on these observations, it may be intuitively assumed that *B. helicrysi* clones are generalist genotypes, fitting the GPG hypothesis. However, the question of putative niche differentiation among aphid superclones, using high-resolution climate data and up-to-date ecological niche analyses, has never been formally addressed.

The aim of this study is to examine whether asexual superclones in the *B. helichrysi* lineage H2 are specialised and differ in their climatic niche, supporting the FNV hypothesis, or if they represent ecological generalists, fitting the GPG hypothesis. To address this question, we used a comprehensive sampling of *B. helichrysi* H2 clones from various locations worldwide. We first characterised their association with endosymbiotic bacteria because the abiotic niche of arthropods can be influenced by endosymbionts. These can both expand the niche space of their hosts by conferring resistance to factors such as temperature stress [26–28], dessiccation [29] or toxic molecules [30], or restrict it, because of their own set of limitations to abiotic conditions such as extreme heat [31,32] or salinity for aquatic organisms [33]. Aphid bacterial endosymbionts are known to modulate many aspects of their hosts’ ecological niches, including thermal tolerance [34]. For instance, it has been shown that two common aphid endosymbionts, *Serratia symbiotica* and *Fukatsuia symbiotica*, can help aphids mitigate the effect of heat stress [26,35–38]. Therefore, it is crucial to identify associations with symbionts when investigating the ecological niches of aphid clonal lineages. The recurrent association of a clone with a particular endosymbiont could indeed lead to a spurious conclusion where the clone appears as a niche specialist though its niche occupancy is driven by the endosymbiont, not the aphid genotype. We then used climatic variables associated with the occurrence data of six globally distributed clones to perform ecological niche divergence analyses. We formally tested whether all clones occur under similar climatic conditions or whether some clones are restricted to specific conditions, showing little overlap with other clones in their climatic preference. The former would suggest that they are indeed widespread generalists, while the latter would suggest that they are ecological specialists.

## METHODS

### Sampling of clones

During the period 1999-2015, we collected over 700 colonies of *B. helichrysi* on 33 host-plant genera in the Americas, Europe, Africa, Asia, and Australia. A colony corresponded to individuals collected from a single plant or, if very few aphids were present, from two neighbouring plants of the same species. Each sampling locality was georeferenced, and the host-plant species was recorded. Full details on sampling methods and specimen conservation are provided in [20]. Vouchers for these samples (i.e. individuals from the same colonies) can be found in the CBGP arthropod collection (https://doi.org/10.15454/D6XAKL) [39]. Among these *B. helichrysi* samples, we selected a subset of individuals that were assigned to six widely distributed H2 clones, based on genotyping using 14 microsatellite markers [19,20] (namely clones “A,” “B”, “C”, “D”, “F” or “G” following methods and nomenclature described by [19]. We obtained a final dataset of 249 individuals (clone A: 52 records; clone B: 46 records; clone C: 87 records; clone D: 7 records; clone F: 27 records; clone G: 30 records) from 10 countries and 163 sampling localities (Fig. 1; Appendix S1 and S2).

**Figure 1.**
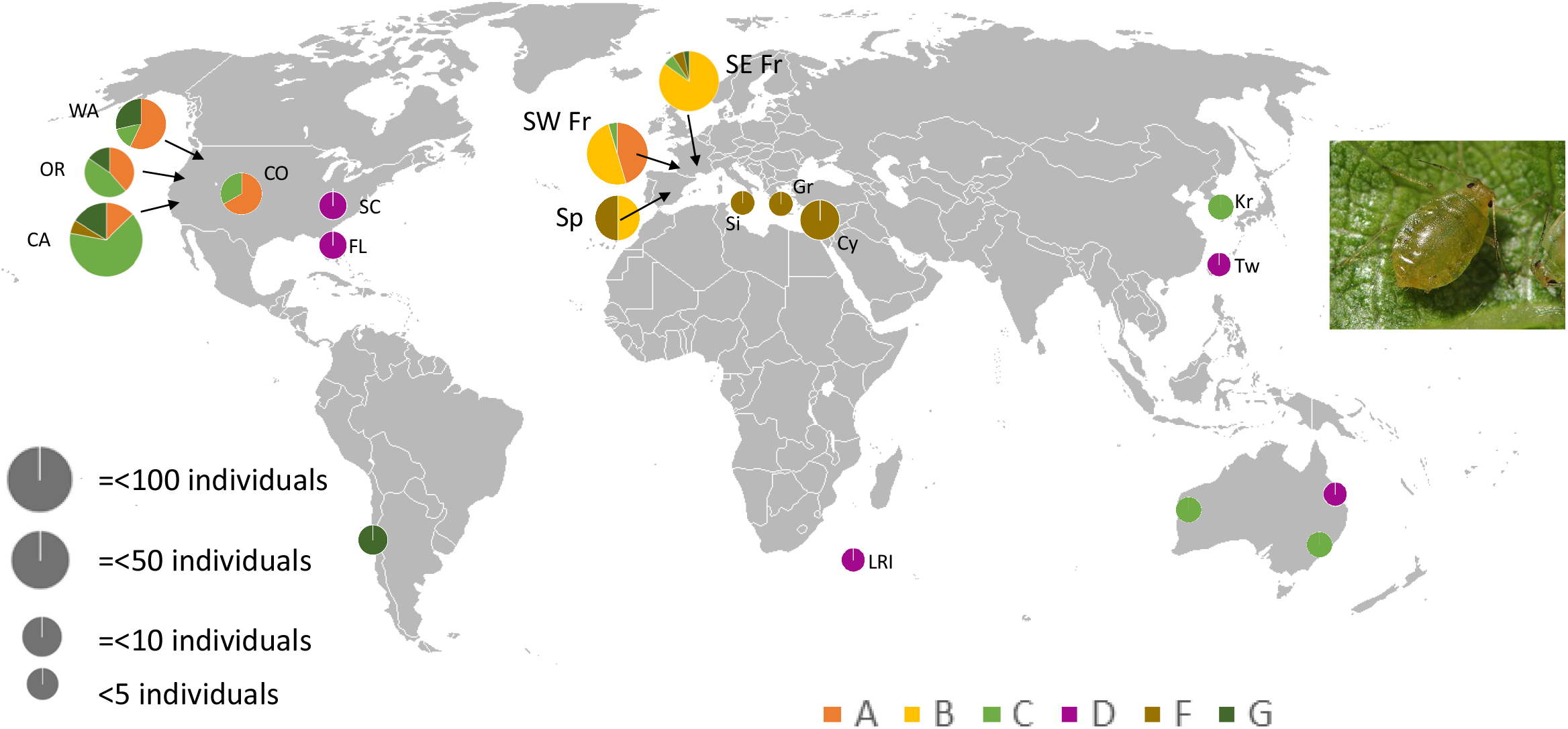
Worldwide distribution of *Brachycaudus helichrysi* superclones. Each clone is represented by a unique color (refer to the color legend at the bottom of the map). The size of each pie chart is proportional to the number of occurrences (see legend on the left side of the map) at each sampling site, and the pie chart slices represent the percentage of occurrences of each clone. Abbreviations: WA: Washington State, OR: Oregon, CA: California, CO: Colorado, SC: South Carolina, FL: Florida, Ch: Chile, SW Fr: South-West France, SE Fr: South-East France, Sp: Spain, Si: Sicily, Gr: Greece, Cy: Cyprus, LRi: La Réunion Island France, Kr: Republic of Korea, Tw: Taiwan, QI: Queensland, NSW: New South Wales

### Characterisation of the endosymbionts associated with each clone

A total of 102 individuals, representative of the six clones and the diversity of their geographic distribution were used for characterising bacterial endosymbionts (Appendix S3). Using DNA extracts from the same individuals utilised for microsatellite genotyping in [19] or [20], we amplified a 251 bp portion of the V4 region of the 16S rRNA gene [40] and used targeted sequencing of indexed bacterial fragments on a MiSeq (Illumina) platform [41] following the protocol described in [42]. Each DNA sample was amplified twice (replicates were conducted on distinct 96-well microplates). We also used negative controls (DNA extraction and PCR controls conducted on blank templates) to filter out bacterial contamination during laboratory procedures. A total of 216 PCR products (comprising DNA extracts and controls) were obtained, pooled, and then separated by gel electrophoresis. Bands based on the expected size of the PCR products were excised from the gel, purified with a PCR clean-up and gel extraction kit (Macherey-Nagel), and quantified with the Kapa Library Quantification Kit (Kapa Biosystems). Paired-end sequencing of the DNA pool was carried out on a MISEQ (Illumina) FLOWCELL with a 500-cycle Reagent Kit v2 (Illumina). Raw sequencing data is available in Dryad repository (https://doi.org/10.5061/dryad.37pvmcvr8).

We first applied sequence filtering criteria following Illumina’s quality control procedure. We then merged paired sequences into contigs with FLASH V.1.2.11 [43] and trimmed primers with CUTADAPT v.1.9.1 [44]. We then used the FROGS pipeline [45] to generate an abundance table of symbiont lineages across samples. In brief, we first filtered out sequences > 261 bp and < 241 bp, then we clustered variants into operational taxonomic units (OTUs) with SWARM [46] using a maximum aggregation distance of three. We identified and removed chimeric variants with VSEARCH [47].

Taxonomic assignment of OTUs was carried out using RDPtools and Blast [48] against the Silva138-16s database (https://www.arb-silva.de) as implemented in FROGS. The resulting abundance table is available in Appendix S3. From the abundance table of OTUs across samples, we transformed read numbers per aphid sample into frequencies (percentages); sequences accounting for < 0.5 % of all the reads for a given sample were excluded [42]. We then only kept OTUs that were present in both PCR replicates of the same sample. All filters resulted in excluding reads found in low abundance that could represent sequencing errors and which were also often found in the negative controls.

Altogether, we found a very low diversity of endosymbiotic bacteria associated with aphid specimens (see full details in the Results section; Appendix S3 & S4). To address if endosymbiotic bacteria communities are different among asexual lineages, we focused further analyses on *Hamiltonella defensa* associated samples because this was the only facultative endosymbiont found in a relatively large number of samples (see the Results section). We fitted a GLM (binomial family) to explore if the probability of finding *Ha. defensa* in sampled aphid specimens depended on clone identity (response variable: presence/absence of *Ha. defensa* in a sample; explanatory variable: clone identity).

### Realised climatic niche divergence

#### Distributional data and bioclimatic descriptors

We investigated realised climatic niche divergence among *B. helichrysi* clones by extracting values of 20 ecologically relevant bioclimatic variables from each sampling locality. Bioclimatic data were obtained from the CHELSA database at a resolution of 30 arc-seconds [49]. These bioclimatic data represent worldwide historical trends in temperature, solar radiation, and precipitation for the period 1979-2013 (Appendix S5). For each clone, we removed duplicated records, i.e., we only allowed one occurrence per pixel of the bioclimatic rasters. After removing duplicated records, the presence dataset encompassed 176 occurrences (Table 1).

**Table 1:** Number of occurrences included in the analyses after removing duplicated occurrences and results of the climatic niche equivalency tests derived from both between-class analyses (i.e., BCA#1: clone D versus all other clones; BCA#2: each clone compared to other clones excluding the clone D)

| Clone | Number of occurrences | Observed Schoener's D index | Niche equivalency test p values |  | Analysis |
| --- | --- | --- | --- | --- | --- |
| Clone A | 34 | 0.52 | 0.005 |  | BCA #2 |
| Clone B | 35 | 0.14 | 0.001 |  | BCA #2 |
| Clone C | 56 | 0.53 | 0.001 |  | BCA #2 |
| Clone D | 6 | 0.06 | 0.052 |  | BCA #1 |
| Clone F | 19 | 0.48 | 0.013 |  | BCA #2 |
| Clone G | 26 | 0.59 | 0.037 |  | BCA #2 |

#### Niche equivalency tests

We employed niche equivalency test to examine the divergence in climatic niches among clones [50]. The analysis was conducted using the *ade4* package [51] and the *ecospat* package [52] in the R environment [53]. We compared the niches of clones in a bidimensional space defined by a between-classes analysis (BCA) [54], calibrated on the entire environmental space represented by climatic values associated with the occurrences of all clones of *B. helichrysi* H2. The BCA - namely the “*BETWEEN-occ*” approach according to [50] - was preferred to other ordination approaches because this technique maximises both inter-individual and between-classes (here, the different clones) variance, aligning with the objective of the study. To assess niche equivalency, we performed subsequent tests comparing the niche of each clone against the collective niche of all other remaining clones, employing niche equivalency test based on Schoener’s index [52,55] We opted for a resolution of 100 X 100 when creating the grid with occurrence densities along gridded environmental gradients. Occurrence data for all clones were then 1,000 times combined and randomly allocated to two datasets of the same size as the dataset under investigation (number of occurrences for the clone being examined versus the number of remaining occurrences). We compared the observed D values between clonal lineages to the distribution of these 1,000 simulated values. If the observed *D* fell within the density of 95% simulated values, the null hypothesis – indicating niche equivalency-could not be rejected.

Outputs of this first BCA (referred to as BCA #1 in the following text) suggested that clone D that occurs in tropical and sub-tropical regions may have a substantially different climatic envelope compared to all other clones, which by contrast occur in temperate regions. Capturing subtle niche differences between most of temperate clones (i.e., all clones except clone D) was thus challenging in this factorial bidimensional BCA-derived environmental space. Indeed, the climatic variables with the highest loading scores on both first axes of the BCA #1 mainly reflect the duality between tropical *versus* temperate climates.

To capture differences between temperate clones, we thus repeated the full procedure while excluding clone D (referred to as BCA #2 in the following text). In this BCA #2 approach, the BCA was calibrated on the entire environmental space represented by climatic values associated with the occurrences of *B. helichrysi* lineage H2, all clones included except clone D.

## RESULTS

### Diversity of endosymbionts associated with B. helichrysi clones

After sequence filtering steps, the high-throughput sequencing of 16S rRNA bacterial genes of 102 aphids and six negative controls resulted in 3 M sequencing reads with an average of 13,680 reads per PCR product. After the application of all necessary filters to remove contaminants, the remaining dataset consisted of 2.7 million sequencing reads, which were clustered into 55 OTUs. This survey revealed an extremely low diversity of bacterial symbionts in *B. helichrysi* clones (Appendix S3 & S4). Most of the reads (75%) were assigned to the aphid primary symbiont, *Buchnera aphidicola*, which was found, as expected, in all samples. Another well-known aphid endosymbiont, *Ha. defensa*, accounted for 3.5% of the reads and was found in 21 out of the 102 aphid specimens. A single sample contained another known aphid endosymbiont, *Erwinia haradae* [56]. Other samples did not host any common aphid endosymbionts. The remaining sequencing reads were either assigned to ubiquitous bacteria (that were sometimes not assigned below family level, such as Chitinophagaceae), or represented bacteria found at very low frequencies, i.e., below 0.5% of the reads in the samples (these are represented in grey in Appendix S4). These bacteria might therefore be biologically irrelevant. Overall, the analysis indicated a scarcity of symbionts associated with *B. helichrysi* individuals, regardless of their clonal lineage.

The endosymbiont *Ha. defensa* was predominantly found in clone C (it was found in 5, 15 and 1 individuals belonging to clones B, C and F respectively (Appendix S3 & S4). According to the logistic GLM, there was a statistically significant effect of clone identity on the presence of the endosymbiont *Ha. defensa* in sampled specimens (NULL deviance: 102.79 on 99 degrees of freedom; Residual deviance: 72.048 on 94 degrees of freedom, χ2 = 30.74, ddl =4, P<0.001).

### Ecological niche divergence between asexual lineages

#### Climatic niche divergence analyses

The results of BCA #1 are presented graphically in Figure 2. The first two factorial axes of the BCA #1 accounted for 61.4, and 20.1 % of the total variation in climate conditions experienced by the clones, respectively. In the BCA #1, the first factorial axis clearly distinguished clone D from all other clones (Fig. 2). Clone D is associated with high annual precipitation (bio12), high amounts of precipitation during the driest and the warmest periods of the year (bio14, bio17, bio18). Clone D is also associated with high temperature during the coldest, warmest and wettest periods of the year (bio5, bio6, bio8), high annual mean temperature (bio1) and low values of mean diurnal range (bio2), isothermality (bio3) and precipitation seasonality (bio15) (Fig. 2). The hypothesis of niche equivalency between clone D and all other clones combined was close to being rejected; Table 1; Appendix S6). The limited sample size of clone D (only six records, albeit from distant sites as shown in Fig. 1) may explain the marginal significance despite the distinct position of clone D in the BCA #1 analysis (Table 1).

**Figure 2.**
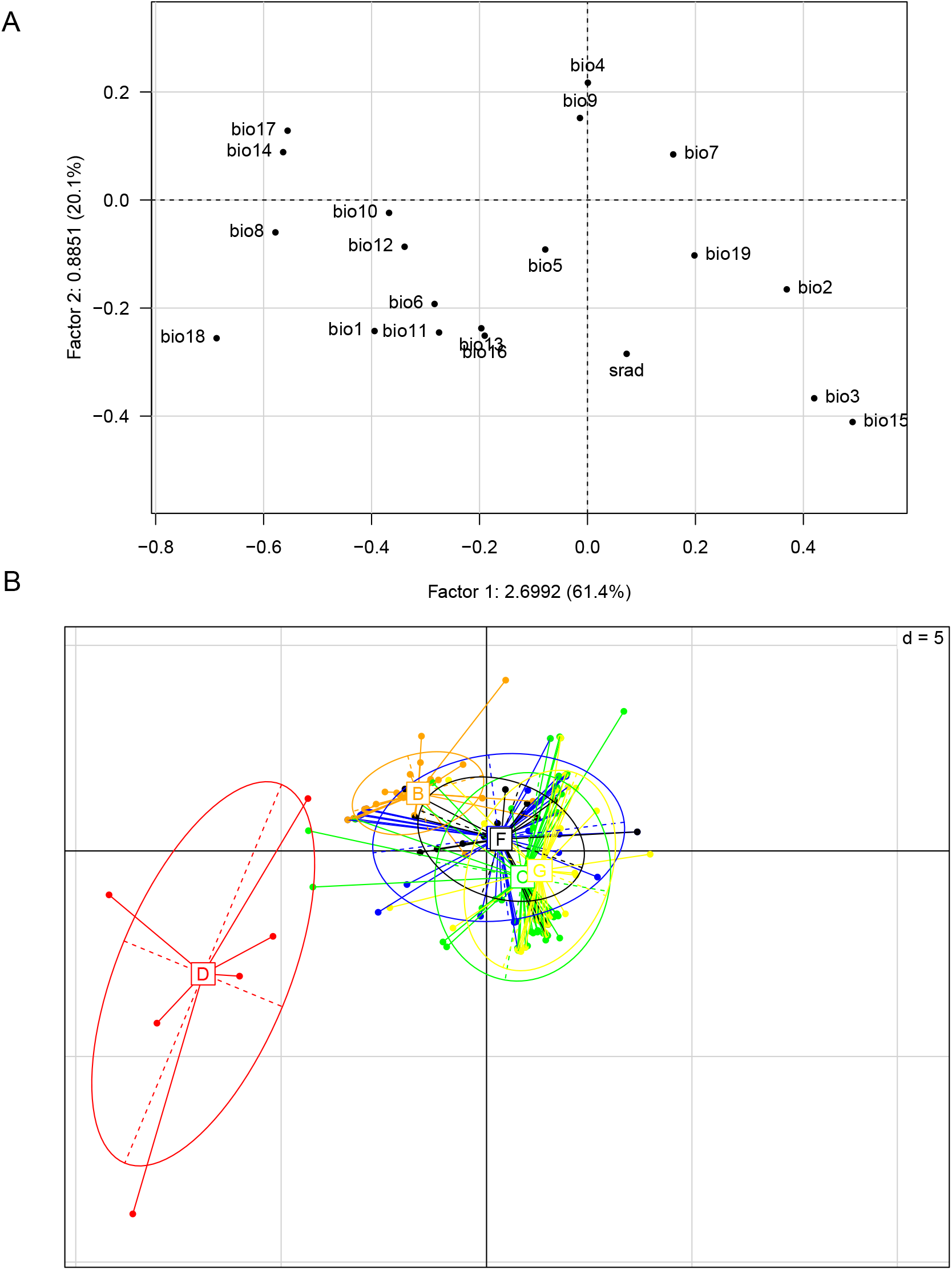
(A) Scores of bioclimatic variables on the first two factorial axes of the between-class analysis #1 (i.e., all clones included). Meaning of abbreviations of climatic variables are available in Appendix S5. (B) Position of six *Brachycaudus helichrysi* clones (i.e., clones A, B, C, D, F and G) along the two first axes of this between-class analysis #1.

Graphical outputs of the BCA #2 calibrated without considering clone D (correlation circles and score plots) are given in Figure 3. The first two BCA #2 axes accounted for 65.2, and 25.3 % of the total variation in climate conditions experienced by the clones, respectively. Based on the BCA #2, the hypothesis of niche equivalency between clones was rejected for each clone (Table 1, Appendix S7). The first BCA #2 factorial axis strongly discriminates clone B from all other clones (Fig. 3). Clone B is associated with relatively high amounts of precipitation during the driest and warmest period of the year (bio14, bio17, bio18), high temperature during the wettest quarter of the year (bio8), low isothermality (bio3) and low rainfall seasonality (bio15) (Fig. 3). Clone A has a positive score on the second BCA #2 factorial axis (Fig. 3). The niche position of clone A is associated with high diurnal range (bio2), low annual solar radiation, a cold winter (bio6, bio11), high annual temperature range (bio7), low annual average temperature (bio1) and high amounts of precipitation during the coldest quarter of the year (bio19) (Fig. 3). Clone C has a negative score on the first BCA #2 factorial axis (Fig. 3). This reflects a niche position associated with relatively low amounts of precipitation during the driest and warmest period of the year (bio14, bio17, bio18), low temperature during the wettest quarter of the year (bio8), high isothermality (bio3) and high rainfall seasonality (bio15) (Fig. 3). Clone F occupies a negative position on the second BCA #2 factorial axis (Fig. 3). The niche position of clone F on the BCA #2 factorial space is associated with low diurnal range (bio2), high solar radiation and warm and dry winter (bio6, bio11, bio19) and high temperature during the wettest quarter of the year (bio8) (Fig. 3). The niche position of clone G is associated with high diurnal range (bio2), isothermality (bio3), and precipitation seasonality (bio15), low temperatures during the wettest period of the year (bio8) and relatively low amounts of precipitations during the driest and warmest period of the year (bio14, bio17, bio18) (Fig. 3).

**Figure 3.**
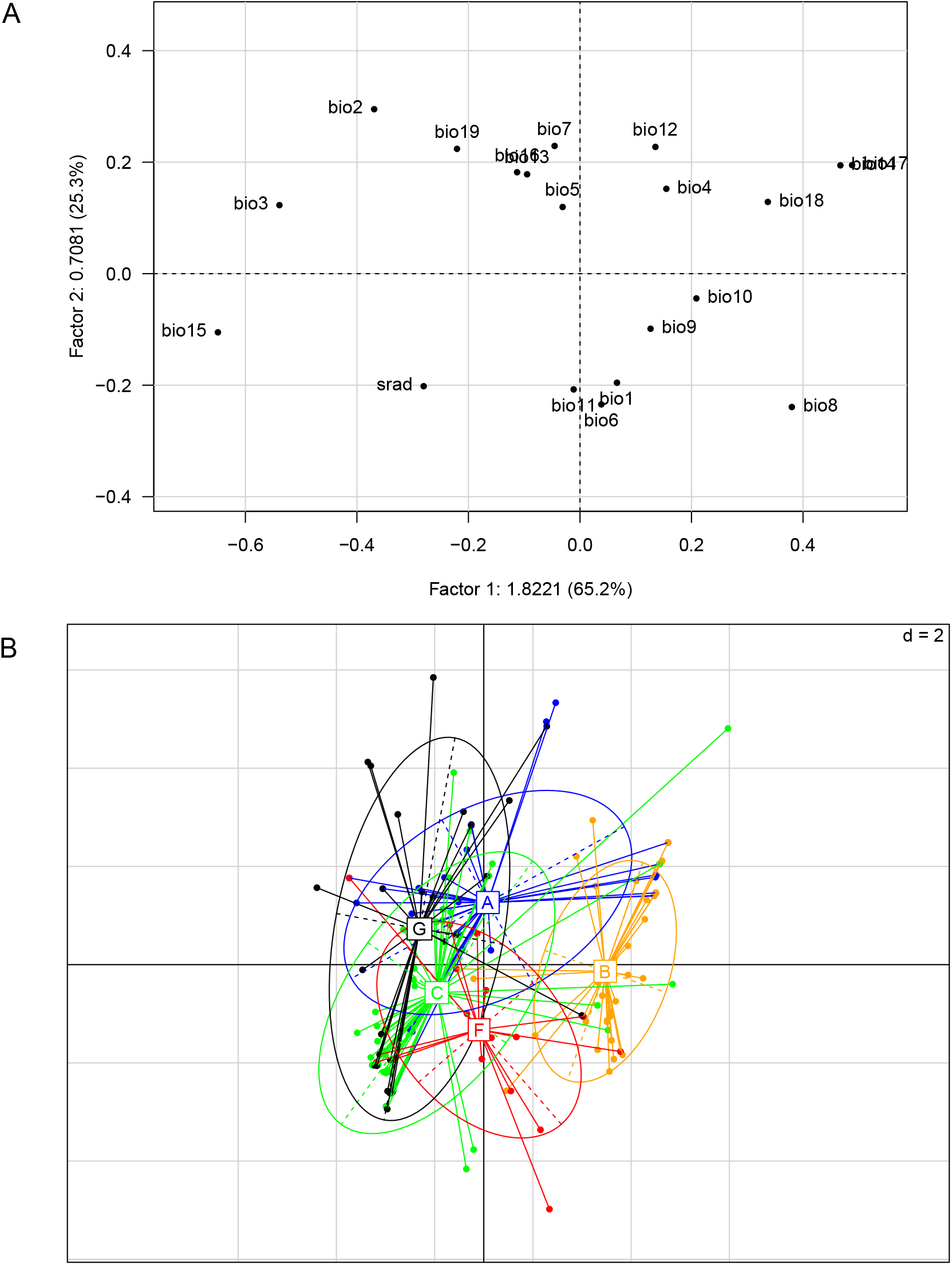
(A) Scores of bioclimatic variables on the first two factorial axes of the between-class analysis #2 (i.e., all clones included except clone D). Meaning of abbreviations of climatic variables are available in Appendix S5. (B) Position of five *Brachycaudus helichrysi* clones (i.e., clones A, B, C, F and G) along the two first axes of this between-class analysis #2.

## DISCUSSION

Understanding the ecological differentiation of asexual lineages provides valuable insights into the maintenance and evolutionary dynamics of asexuality, shedding light on the factors driving clones’ geographic distributions and ecological success. Aphids yield persistent clonal lineages that can expand over large geographic scales, sometimes over continents. Among the aphid species that comprise such lineages are *Myzus persicae* [25], *Sitobion avenae* [57], *Aphis gossypii* [58], *B. helichrysi* [19] and *Melanaphis sacharri* [59]. Because of their wide distribution and apparent ecological success, these so-called superclones are intuitively assumed to be generalist genotypes that show a high degree of phenotypic plasticity and therefore adaptability to various environments [24,57,60]. Taking advantage of a worldwide sampling of *B. helichrysi* and using multivariate ecological niche analyses, we provide here strong evidence against this assumption. We reveal significant differentiation in the realised climatic niche of six *B. helichysi* superclones. Though it was represented by few occurrences, we found potential niche specialisation of one clone (clone D) to tropical or sub-tropical climatic conditions (Fig. 2; Table 1). The remaining clones were found in geographic areas characterised by temperate climatic conditions but showed some specificity within this range (Fig. 3). For instance, clone F was clearly restricted to the Mediterranean-climate regions, while clones A and B showed apparent niche specialisation to colder temperate climates. Hence, despite their incredibly large geographic distributions, *B. helichrysi* superclones better fit the FNV hypothesis than the GPG.

Previous studies had also refuted the assumptions that aphid clonage lineages were ecological generalists. For instance, it has been shown that, in both *Aphis gossypii* and *Melanaphis sacchari*, long-term asexual lineages have distinct sets of preferred host-plants [58,61]. Some clonal lineages of *Acyrtosiphon pisum* that have spread from Europe to South America also appear to be specialised on specific host-plants, but they probably each derive from a host-specialised sexual lineage, which implies that ecological specialisation is not characteristic of the clone itself but rather inherited from a sexual biotype [62]. Studies of *S. avenae* showed that though clonal lineages were highly plastic in their feeding behaviour [63], they could be characterised by different resistance to insecticides [64,65], also suggesting some type of clone niche specialisation. Finally, using experimental approaches, it was showed that *M. persicae* superclones found in Australia did not have a better mean fitness across diverse plants than their sexual counterparts, also casting doubt on a GPG hypothesis for the emergence of these clones [66]. But, to our knowledge, the present study is the first one that presents evidence for aphid superclones’ adaptation to climate conditions. Beyond studies of superclone niche occupancy, this is also the first evidence that within a cosmopolitan aphid species, distinct lineages can have a preferred set of climatic conditions. Aphid “biotypes” are generally designated based on biological characteristics such as different sets of host plants, but, to our knowledge, there are no studies showing that aphid biotypes differ in their climatic tolerance. Gilabert et al. [67] had previously suggested that climatic specialisation of aphid clones was possible, as they showed that two clonal lineages *of Rhopalosiphum padi* exhibited different geographical clines. However, their analysis was restricted to Northern France and did not include any statistical analyses of climatic variables.

The signal of climatic differentiation detected in our study is robust. First, we sampled *B. helichrysi* H2 populations in five continents and across a wide range of climatic conditions, and therefore, we likely have captured well the realised climatic niche of this lineage. Second, we found similar patterns of climatic niche occupation in spatially independent biogeographic areas for several clones. For example, clone D was consistently found under tropical and subtropical climate in four biogeographic regions i.e., Australia, Asia, North America, and Indian Ocean islands (Fig. 1). Similarly, clone F, which mainly occurs in the Mediterranean basin, was also collected in a similar Mediterranean-climate habitat in the United States (i.e., California, Fig. 1). This suggests that the signals of niche specialisation are not due to dispersal limitations of these clones. The disjunct distribution of most clones is striking. It agrees with previous studies that show that some common aphid pests are regularly transported across the globe through human activities [68]. It also legitimates using the entire environmental space defined by all clones’ occurrences as the background region for each clone in niche equivalency tests. The signal of climatic niche specialisation is also unlikely to be due to biotic associations (e.g., association of clones with specific host plants) limiting each clone’s geographic range, as similar biotic conditions are unlikely to be replicated between distant sites. In agreement with this assumption, it was showed that there were no significant differences between host-plant ranges among clones of *B. helichrysi* [19]. Finally, we found a very low diversity of facultative bacterial endosymbionts in sampled specimens. The endosymbiont *S. symbiotica* that is known to play a role in aphids resistance to heat stress [26,38] was not found in any of our samples, not even in samples of clone F, which is specifically found in areas characterised by hot summers. Furthermore, apart from the preferred association of *Ha. defensa* with samples from clone C, our results reveals that asexual clones of *B. helichrysi* do not shelter different endosymbionts. Hence, altogether this suggests that associations with facultative endosymbiont do not play a role in clone niche occupancy. The only recurrent association found in our study was the one of clone C with *Ha. defensa*. This symbiont is known to induce resistance to parasitoids, but this protection diminishes at elevated temperatures [69]. Therefore, it is very unlikely that it is *Ha. defensa* that drives clone C climatic niche occupancy. Elucidating why *Ha. defensa* is predominantly found in clone C but not in other clones is beyond the scope of this paper. Nevertheless, given the literature on this endosymbiont, we can hypothesise that its association with clone C might be favoured because this clone occurs in regions exhibiting very even temperatures throughout the year (high isothermality) which might enhance the protective benefit of *Ha. defensa*.

As stated in the introduction, the GPG and the FNV have both been supported by empirical observations and experimental approaches in different organisms [10]. Since then, among arthropod studies FNV has been evidenced in *Timema* stick insects, where clonal lineages show narrower realised niches than their sexual relatives [70], and water fleas, where clonal composition of populations is structured by environmental variables [71–73]. On the other hand, arguments supporting the GPG have been found in ambrosia beetles, where two clonal lineages show very similar niche occupancy in terms of host plant utilisation [74] and asexual *Daphnia*, where the invasion and dominance of a single clone has been observed in several lakes throughout a wide geographic range and broad environmental conditions [75]. Among empirical approaches that compare the geographical distribution of different clonal lineages, few have employed statistical investigations of climatic factors underlying these distributions. Coughlan et al. [76] found clonal lineages in hawthorns with large ecological tolerances, consistent with the GPG hypothesis, while Greenwal et al. [77] found significant climatic niche differentiation among unisexual lineages of *Ambystoma* salamanders. But in many species that encompass asexual lineages, when these lineages expand over large geographic areas, they are often assumed to fit the GPG hypothesis without further analyses of environmental variables. This is the case in orabid mites [78] where geographic range sizes have been explicitly used as a proxy for specialisation and a GPG conclusion has been drawn based solely on range size comparison of sexual and asexual populations. By capturing replicated climatic conditions across the globe, we show here, that within the large climatic envelope that can sustain long-term asexuality in aphids (i.e., overall low seasonality with mild or warm winters), there are finer variations to which clonal genotypes might be adapted. Before concluding that climatic adaptation is completely endogenous (i.e., the product of the aphid genome), it might also be interesting to investigate the *Buchnera* strains that are associated with *B. helichrysi* clonal lineages. Indeed, it has been demonstrated that the obligate symbiont *B. aphidicola* can differ in a gene coding for a heat-shock protein [79], which then helps aphids resist temperature stress. Genetic variations in *Buchnera* might therefore also play a role in determining abiotic niche occupancy.

## Conclusions

The present study provides evidence for intraspecific climatic niche divergence in a cosmopolitan aphid species, which is not mediated by associations with facultative endosymbionts. It suggests that the high degree of ecological specialisation in *B. helichrysi* clones shapes the geographic areas they can occupy. To our knowledge, climatic adaptation has seldom been hypothesised to underlie the spatial distribution of intraspecific genetic diversity in aphids, the host plants having been seen as the main factors explaining their phylogeographic lineage distributions [58,61,62,80]. This study adds to the evidence that aphid lineages can have their own set of climatic limitations beyond the availability of suitable host plants [81]. Despite the strong signal observed in our study, correlative distribution-based ecological niche analyses must always be interpreted guardedly [82] since they only depict the realised niche, i.e., a subset of the full range of environmental conditions suitable for a taxa [83]. Therefore, the adaptation of *B. helichrysi* superclones to climatic conditions need to be confirmed with experimental approaches manipulating temperature and humidity in controlled laboratory conditions. Altogether the presence of some clones in more than three continents, despite their high degree of specialisation, is a testimony of the intensity of human-mediated long-distance dispersal events at a global scale. If clonal genotypes were ecological generalists, we could expect that they have expanded by natural dispersal surviving along the way through non-preferred environments. Our results rather suggest repeated multiple human-mediated long-dispersal introduction events and subsequent environmental filtering (i.e., local selection of climatically-adapted clones).

## ACKNOWLEDGEMENTS

We thank Joséphine Piffaretti and Armelle Coeur d’acier for help in various aspects of this study.

## Data availability statement

Information on sampling are available in the database PhylAphidb@se. Raw data are available on the repository Dryad: https://doi.org/10.5061/dryad.37pvmcvr8.

