## Supplementary material for "Asexual lineages of a cosmopolitan pest aphid are associated with different climatic niches": Appendix S2 (6-page pdf file).pdf

**Superclone A − North America**

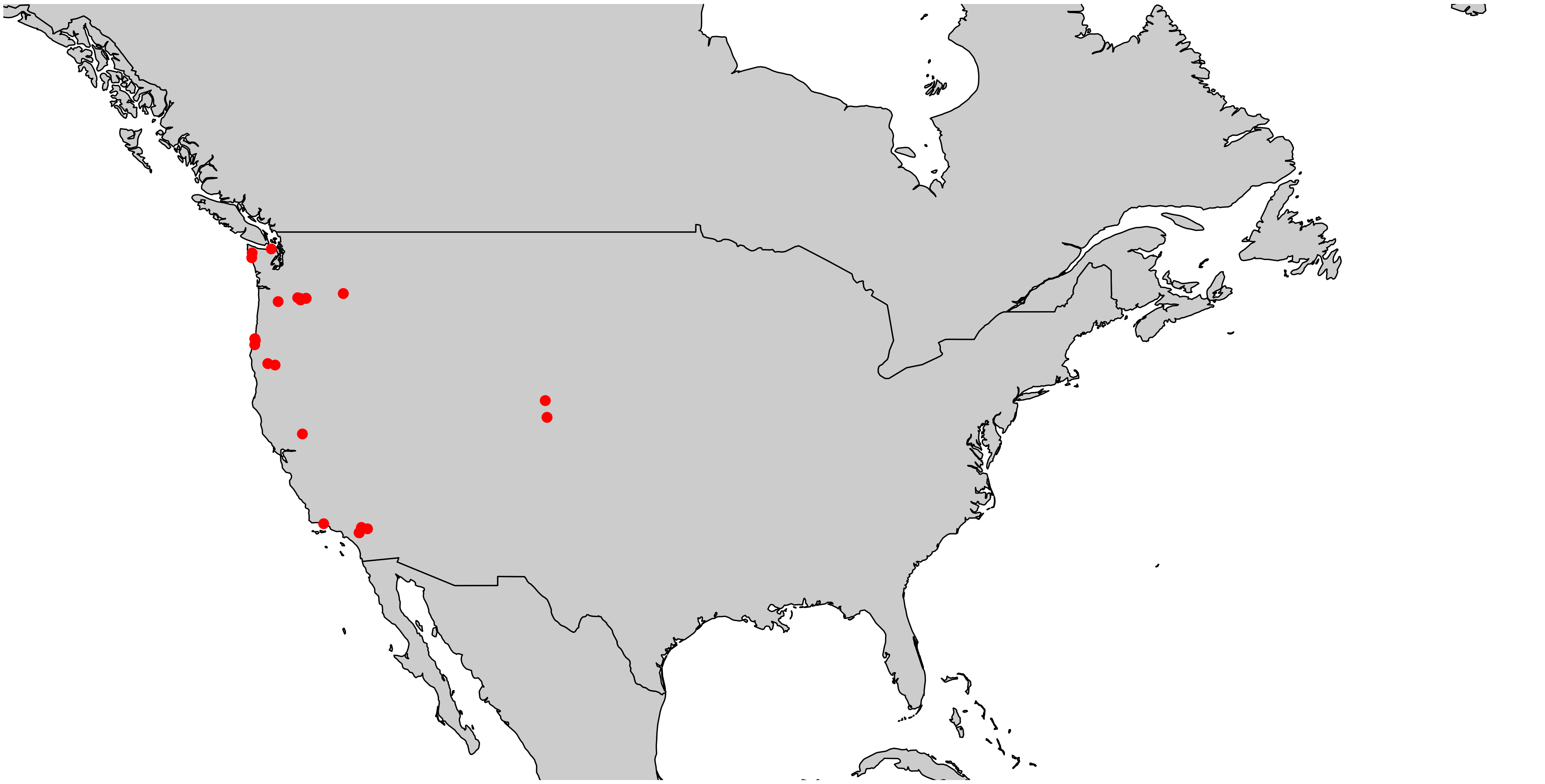

**Superclone A − Europe**

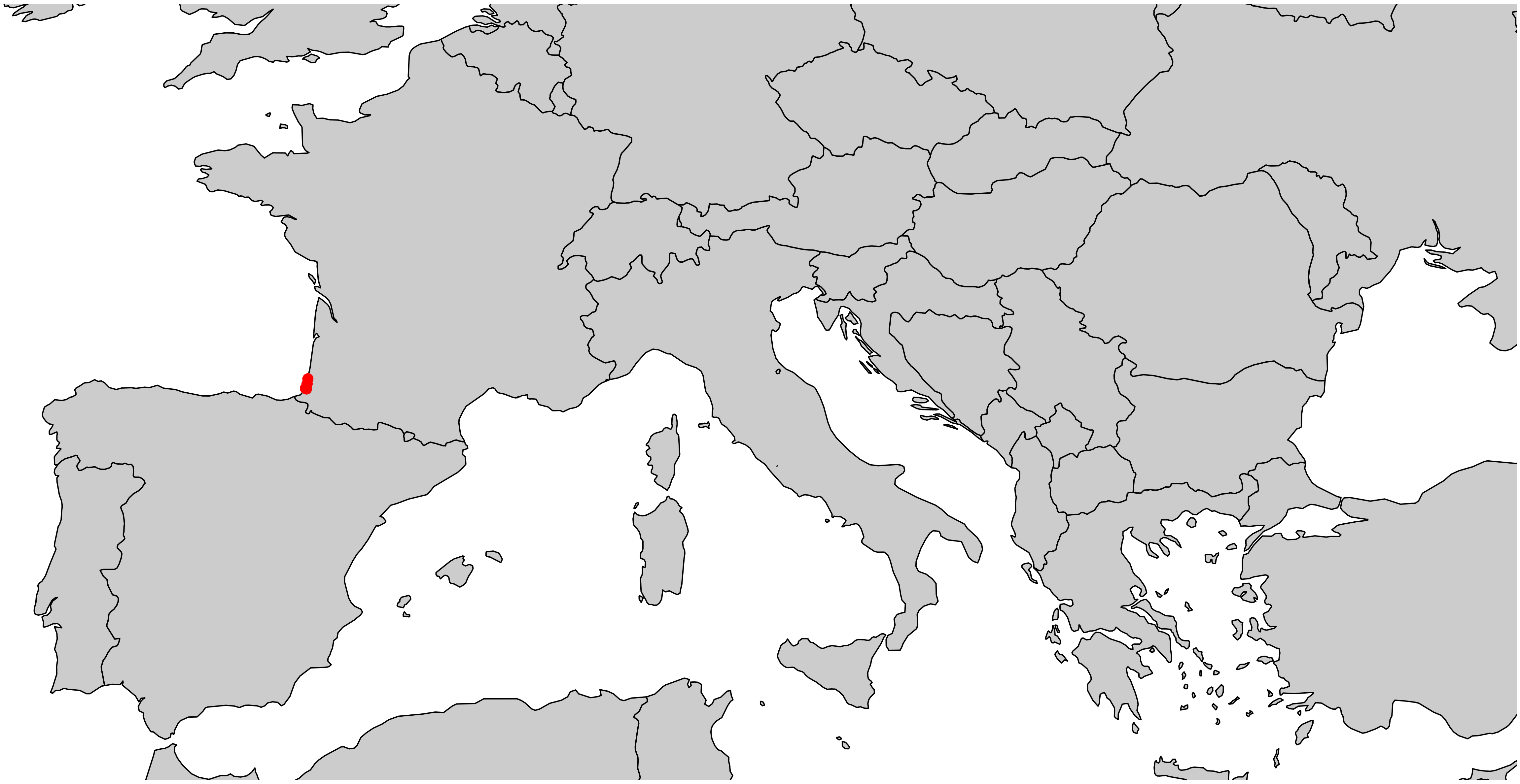

### Superclone B − Europe

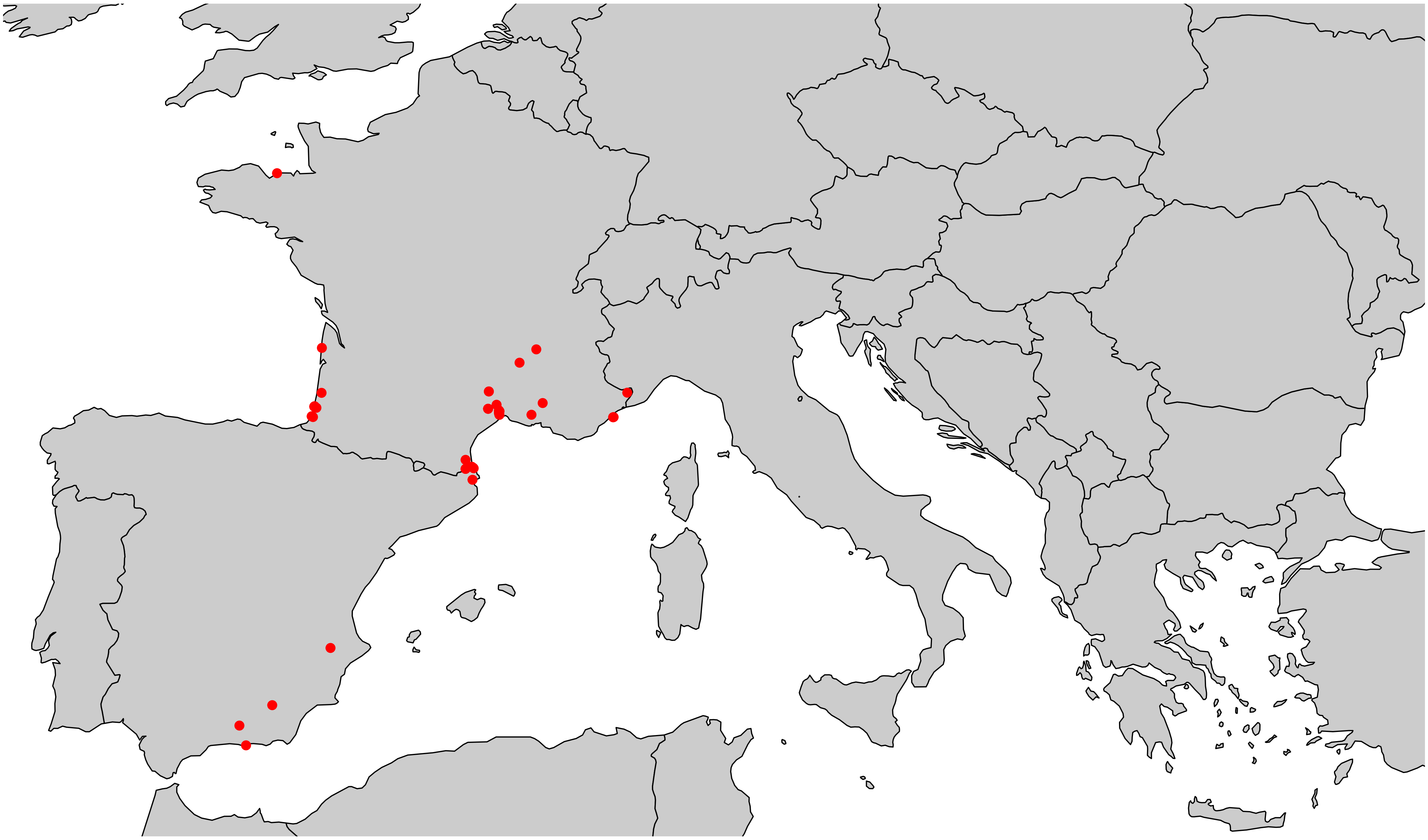

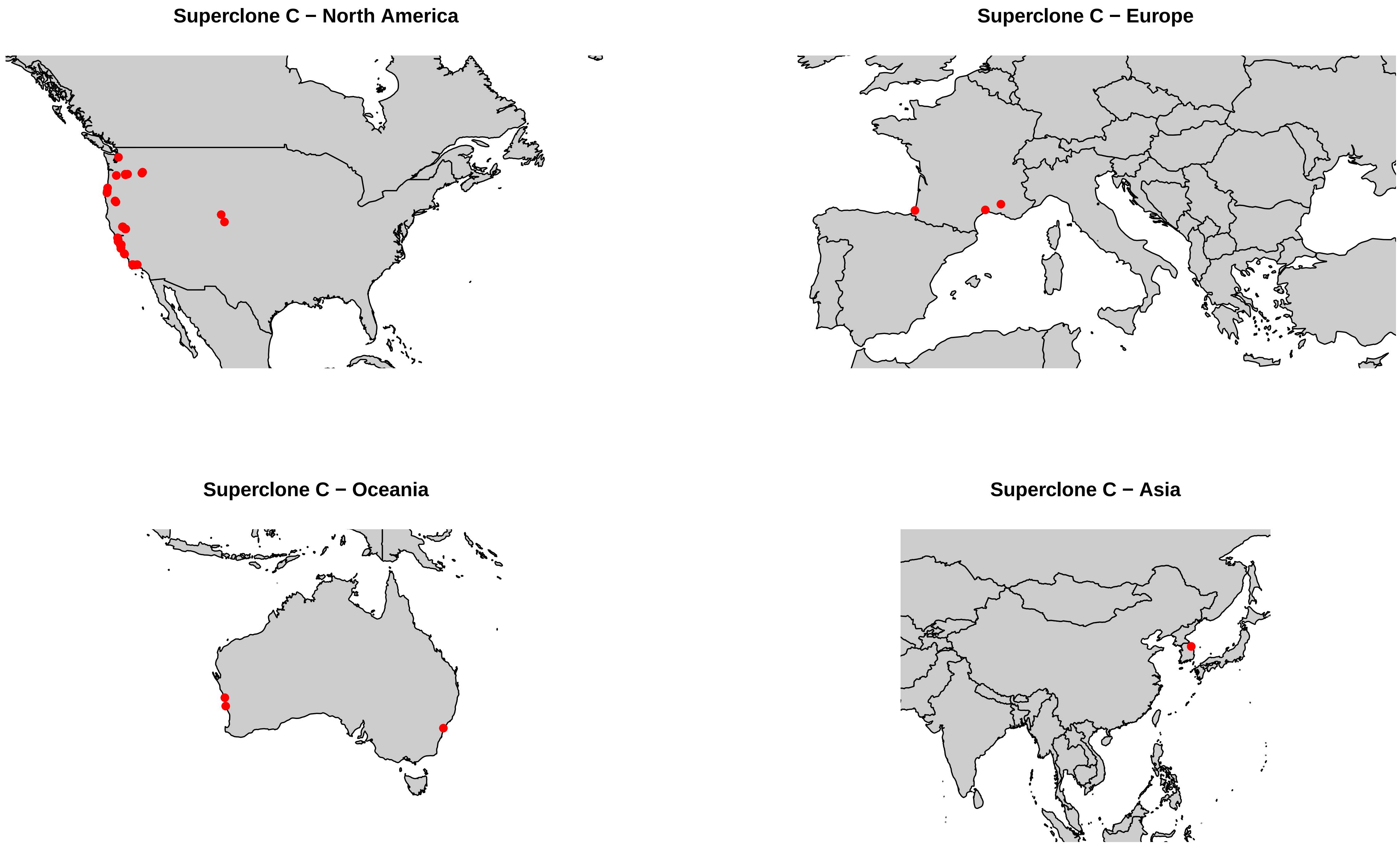
Superclone C − North America
Superclone C − Europe
Superclone C − Oceania
Superclone C − Asia

Superclone D − North America

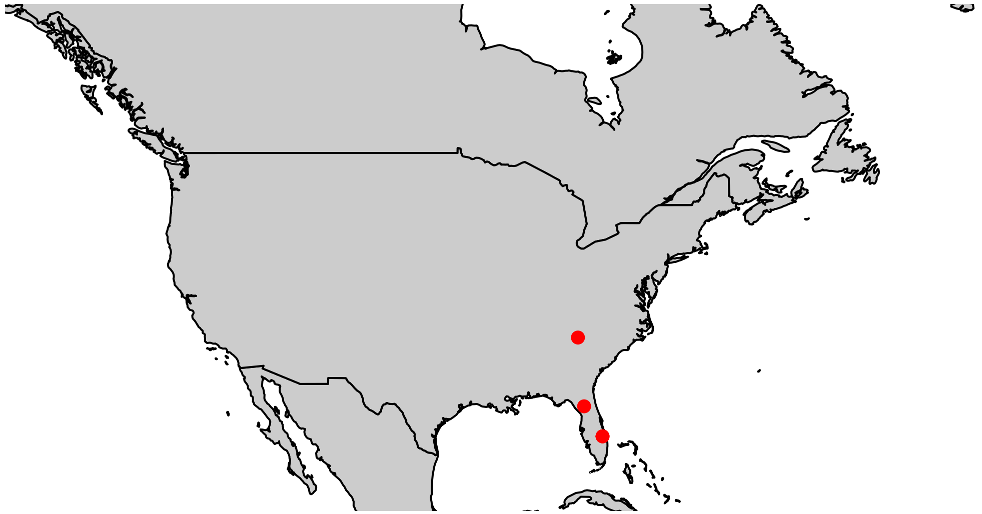

Superclone D − Asia

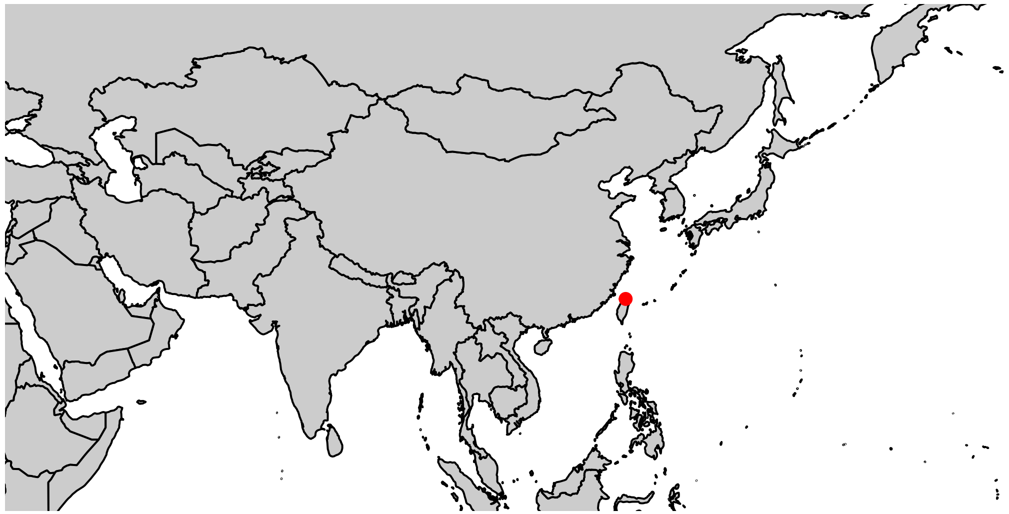

Superclone D − Oceania +  Indian ocean islands

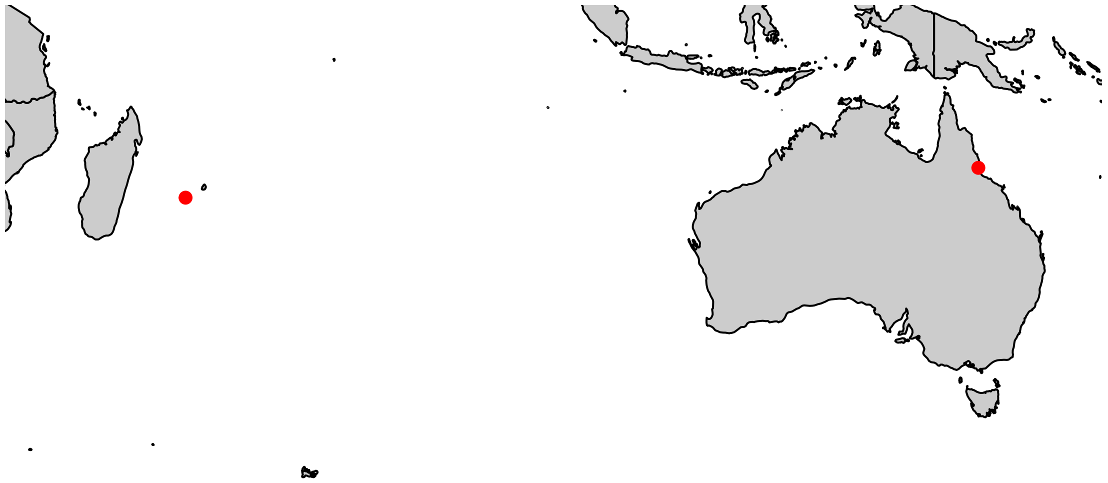

**Superclone F − North America**

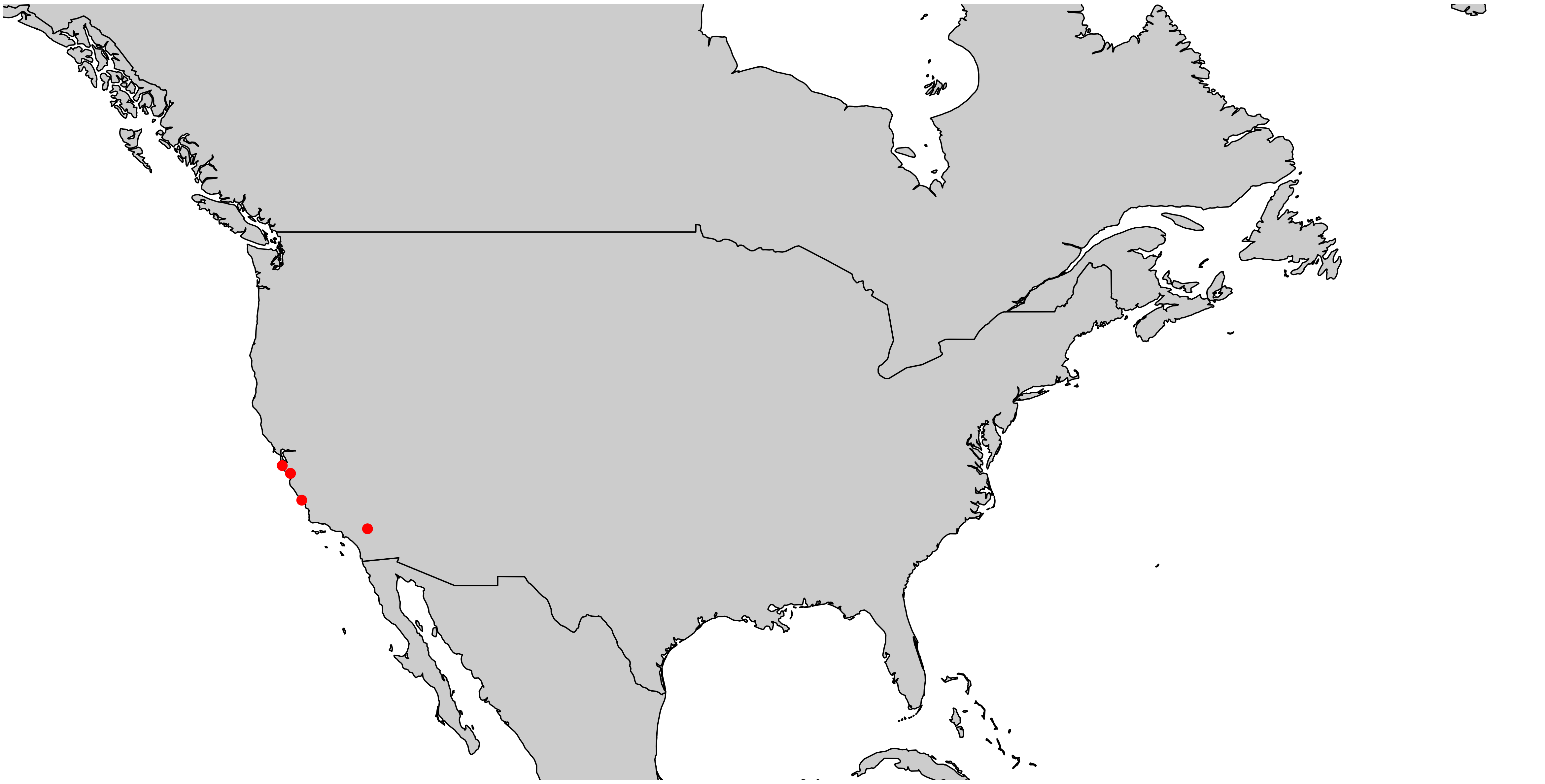

**Superclone F − Europe**

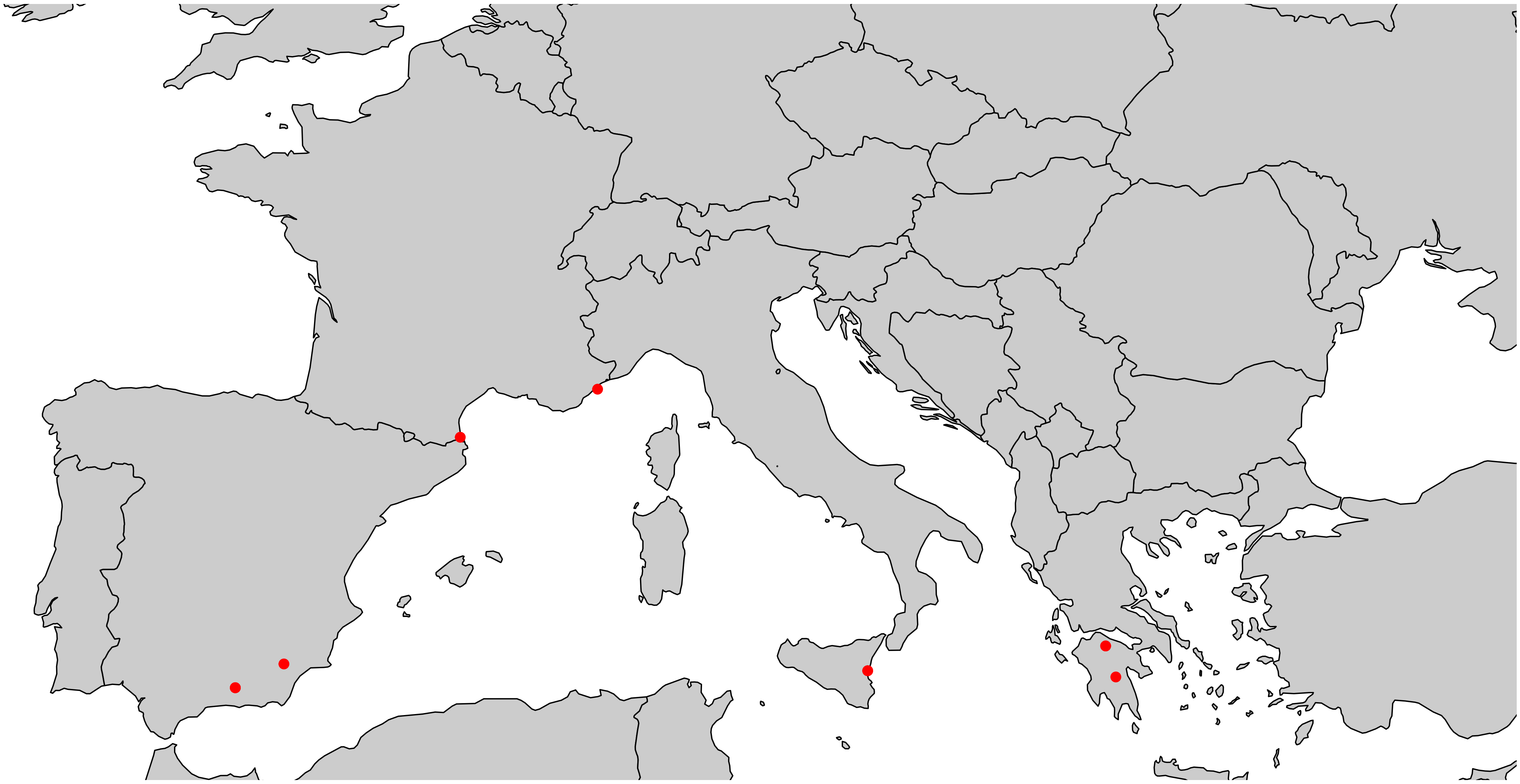

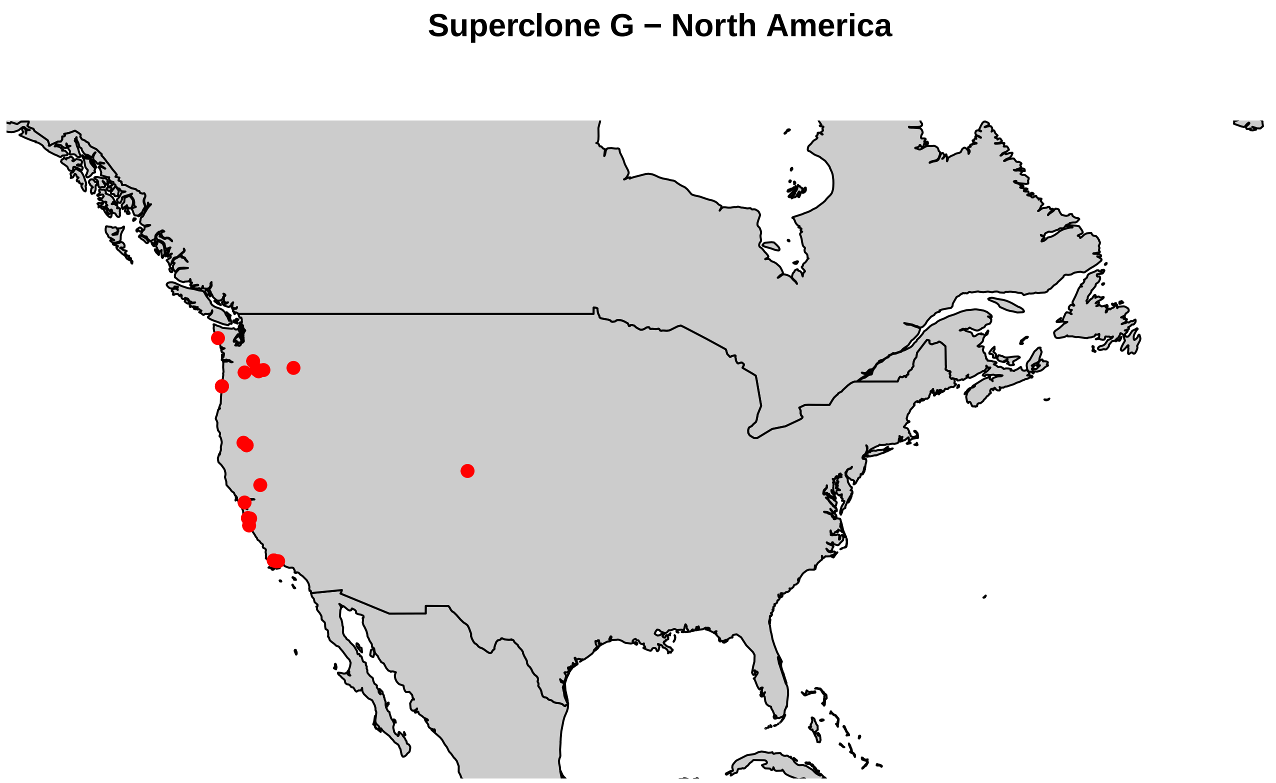
Superclone G − North America

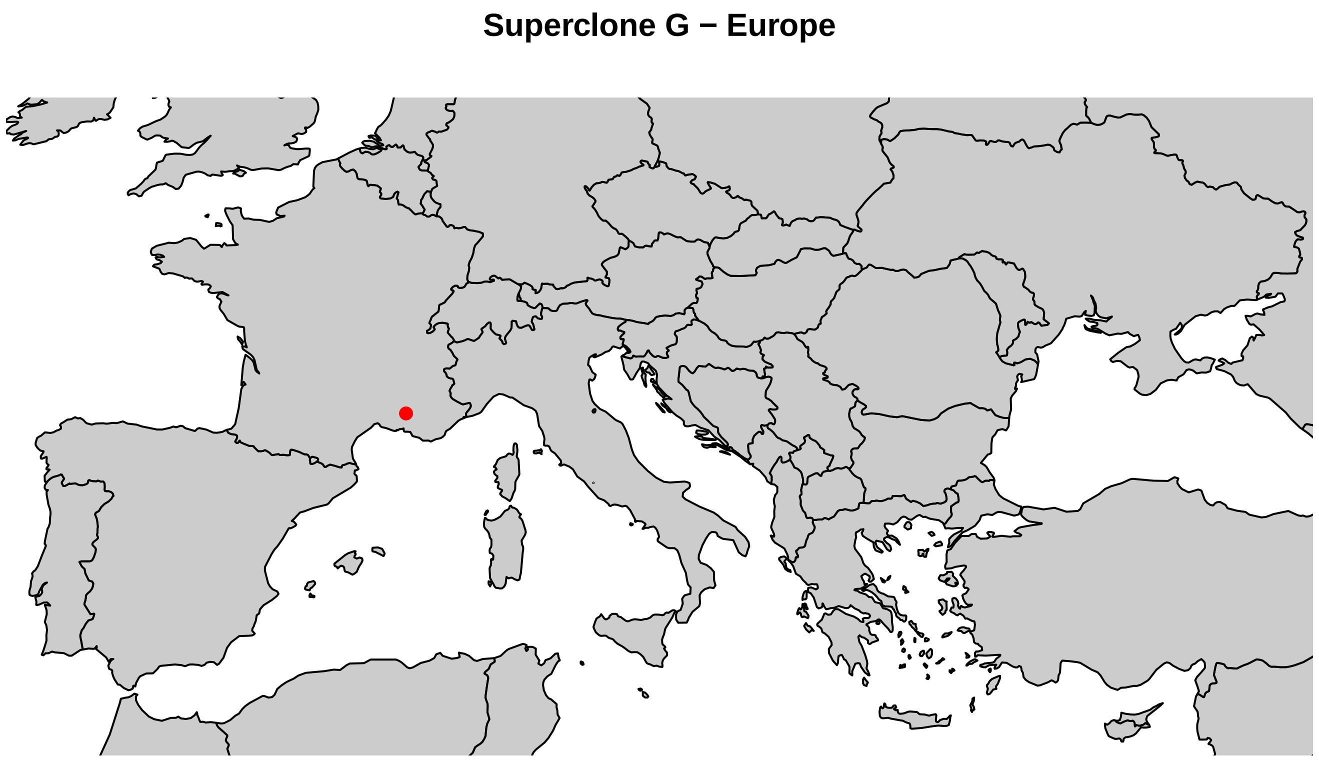
Superclone G − Europe

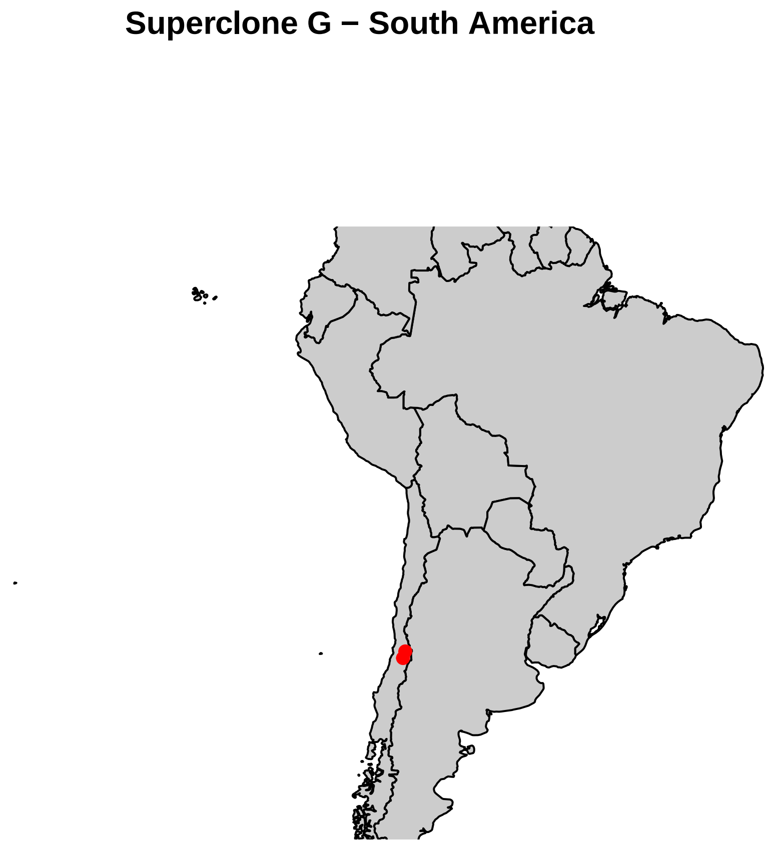
Superclone G − South America
