## Supplementary material for "Asexual lineages of a cosmopolitan pest aphid are associated with different climatic niches": Appendix S5 (one doc file).docx

| Bio1 | Annual mean temperature |
| --- | --- |
| Bio2 | Mean diurnal range (mean of monthly (max temp - min temp)) |
| Bio3 | Isothermality (Bio2/Bio7) (×100) |
| Bio4 | Temperature seasonality (standard deviation ×100) |
| Bio5 | Max temperature of the warmest month of the year |
| Bio6 | Min temperature of coldest month of the year |
| Bio7 | Temperature annual range (BIO5-BIO6) |
| Bio8 | Mean temperature of wettest quarter of the year |
| Bio9 | Mean temperature of driest quarter of the year |
| Bio10 | Mean temperature of warmest quarter of the year |
| Bio11 | Mean temperature of coldest quarter of the year |
| Bio12 | Annual precipitation |
| Bio13 | Precipitation of wettest month of the year |
| Bio14 | Precipitation of driest month of the year |
| Bio15 | Precipitation seasonality (coefficient of Variation) |
| Bio16 | Precipitation of wettest quarter of the year |
| Bio17 | Precipitation of driest quarter of the year |
| Bio18 | Precipitation of warmest quarter of the year |
| Bio19 | Precipitation of coldest quarter of the year |
| Srad | Downward total solar radiation at the surface accumulated over the year |
