## Supplementary material for "Asexual lineages of a cosmopolitan pest aphid are associated with different climatic niches": Supplementary material captions.docx

**Appendix S1** Sampling details on presence records collected for six *Brachycaudus helichrysi* clones (i.e., clones A, B, C, D, F and G).

**Appendix S2** Global mapping of occurrence data collected for six *Brachycaudus helichrysi* clones (i.e. clones A, B, C, D, F and G). Occurrence data is represented by a red circle on the maps.

**Appendix S3** Results of endosymbiotic bacteria genotyping.

**Appendix S4** Distribution of major bacterial associates across samples of *Brachycaudus helichrysi* using 16SS V4 rRNA classification of amplicon sequence variants.

**Appendix S5** List of the twenty bioclimatic variables taken into account in the niche equivalence analyses. Bioclimatic data were obtained from the CHELSA database at a resolution of 30 arc seconds.

**Appendix S6** Results of the niche equivalency test comparing clone D to all other clones. The graphs at the top of the figure represent the realised niche of clone D (top left) and all other clones (top right) on the first two axes of between-class analysis #1 (BCA#1). The solid and dotted lines represent respectively 100% and 50% of the background environment on which the BCA was performed. The grey shading represents the smoothed clone occurrence density. The correlation circle (bottom left) represents the scores of bioclimatic variables on the first two factorial axes of BCA #1. The histogram (bottom right) shows the observed Schoener’s D index of niche overlap (i.e., the bar with a red diamond) and the simulated niche overlap values.

**Appendix S7** Results of the niche equivalency test comparing each clone to all other clones (except clone D). The graphs at the top of the figure represent the realised niche of a clone (top left) and all other clones (top right) on the first two axes of between-class analysis #2 (BCA#1). The solid and dotted lines represent respectively 100% and 50% of the background environment on which the BCA was performed. The grey shading represents the smoothed clone occurrence density. The correlation circle (bottom left) represents the scores of bioclimatic variables on the first two factorial axes of BCA #2. The histogram (bottom right) shows the observed Schoener’s D index of niche overlap (i.e., the bar with a red diamond) and the simulated niche overlap values.
