## Supplementary figures and images for "Asexual lineages of a cosmopolitan pest aphid are associated with different climatic niches"

### Appendix S4 (one png file).png

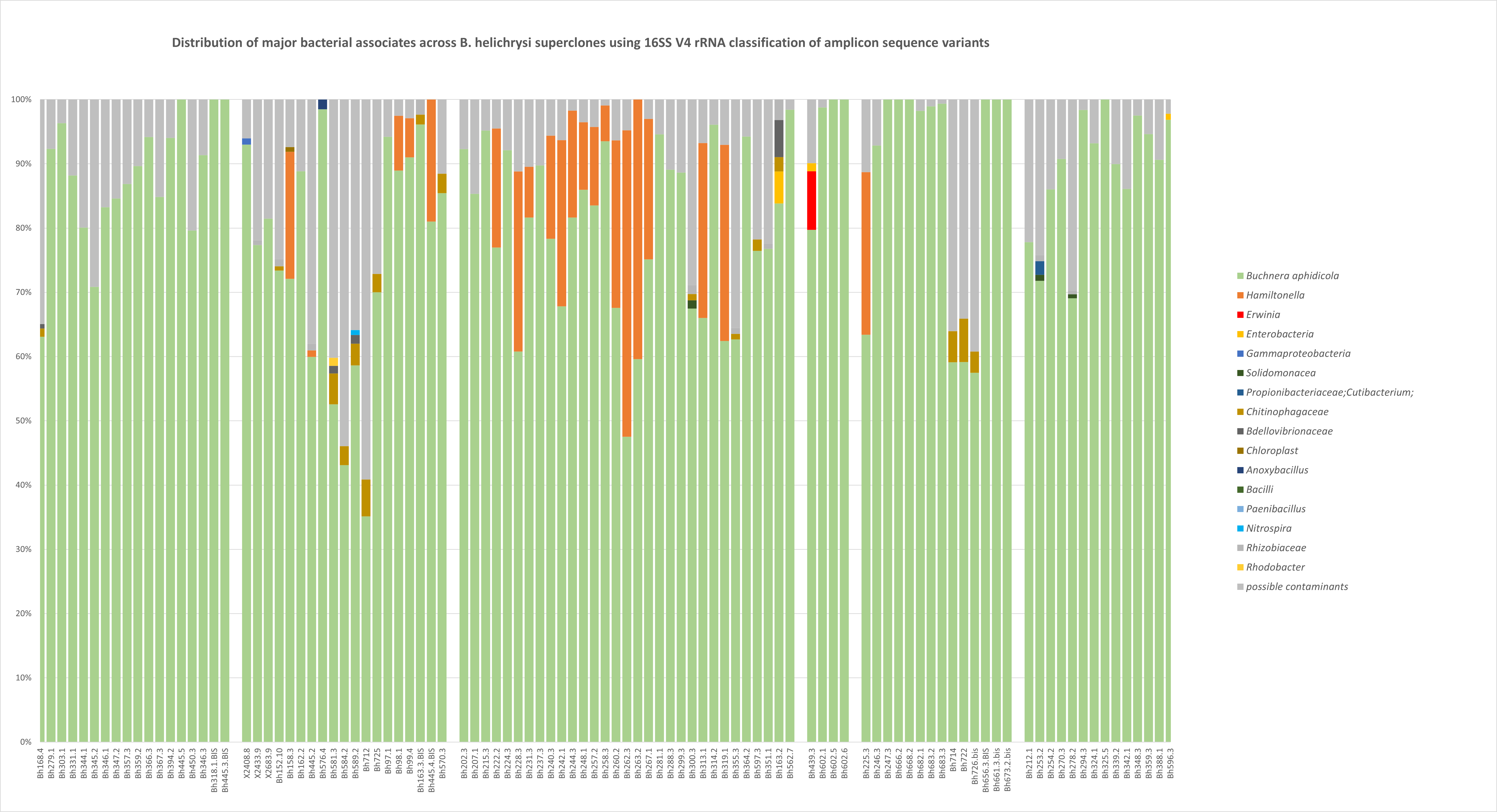

### appendix_S6 (one pdf file).pdf

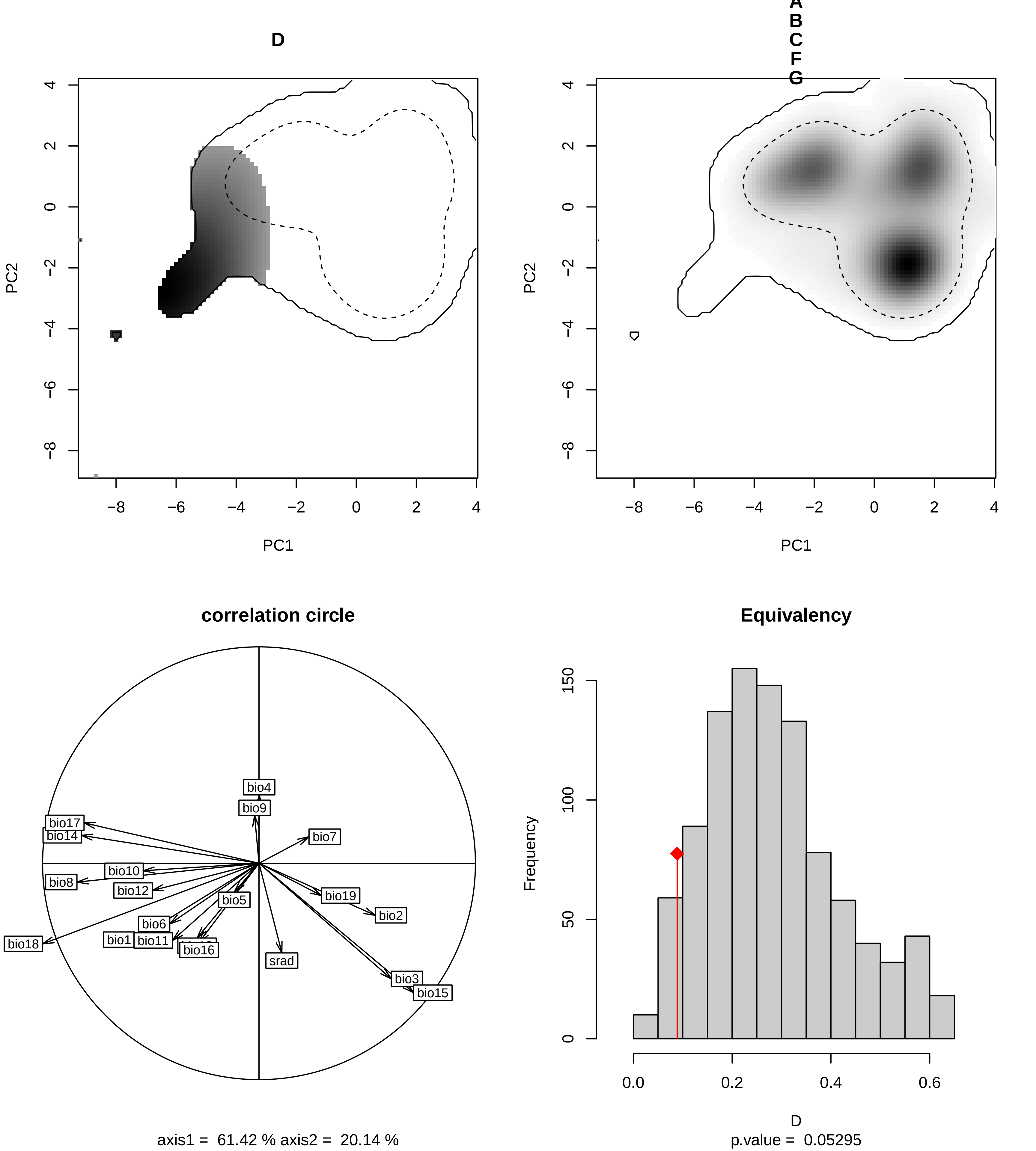
